# BraiNN: A Modern Simulator for Clinically Feasible Personalized Whole-Brain Network Modeling

**DOI:** 10.64898/2026.07.08.737156

**Authors:** Alessandro Fasse, Chiara Billi, Victor Garvalov, Matthew Morvan, Taylor Newton, Niels Kuster, Esra Neufeld

## Abstract

Personalized whole-brain modeling aims to transform treatment planning for neurological disorders by enabling patient-specific simulations of brain network dynamics. Neural mass models (NMMs) offer a tractable compromise between biophysical detail and computational cost and can be directly linked to macroscopic observables such as EEG. However, scaling NMMs to whole-brain networks with realistic connectivity, conduction delays, and cortical surface resolution—and fitting them to individual patient data—imposes computational demands that existing frameworks cannot meet at clinically relevant timescales. Here we introduce *BraiNN*, a JAX-based Python framework for large-scale neural mass modeling that achieves speedups of up to two to three orders of magnitude over existing tools by leveraging GPU/TPU-accelerated, XLA-compiled array computation. BraiNN combines a region-level Jansen–Rit network with a subject-specific cortical surface mesh of coupled neural mass models and biophysically grounded EEG forward modeling via reciprocity-based lead fields. Its fully differentiable computational graph enables a hybrid personalization pipeline that pairs Bayesian optimization for global parameter exploration with gradient-based refinement, completing EEG-driven spectral fitting of an eight-dimensional parameter space in approximately 2–3 hours on a single consumer GPU—compared to multiple days with conventional neural mass modeling software. Numerical verification against established benchmarks confirms that BraiNN faithfully reproduces canonical synchronization and bifurcation dynamics of Jansen–Rit networks.

By reducing the time requirements for personalizing a high-detail whole-brain surface model from days to a few hours on consumer-grade hardware, BraiNN brings personalized brain network modeling closer to practical use in clinical contexts. We anticipate that BraiNN will serve as a foundation for patient-specific digital twins and EEG-guided neuromodulation planning.

## Introduction

Over the last decade, noninvasive neuromodulation and advanced neuroimaging have started to move from population-based protocols toward individualized interventions. Techniques such as navigated transcranial magnetic stimulation increasingly rely on subject-specific structural and functional maps to define stimulation targets and to interpret downstream effects. Recent work on individualized brain mapping and parcellation demonstrates that inter-individual variability in cortical organization is substantial and clinically relevant, and that group-level atlases may obscure precise differences that matter for neuromodulation planning (1, 2). This creates a demand for modeling frameworks that can exploit personalized anatomical and functional data while remaining practical within clinical and translational workflows.

At the same time, it is well recognized that brain dynamics operate across multiple spatial and temporal scales, and that no single modeling level can fully capture this complexity. Large-scale neuron and synapse–based simulations provide a principled route to modeling and have been proposed as tools to understand both normal and pathological network states (3). However, even with modern high-performance computing, such bottom-up simulations remain technically demanding and computationally expensive, particularly when repeated simulations are required for parameter exploration, model fitting, or patient-specific scenario testing. Conceptual and methodological issues further complicate their straightforward application to individualized, time-constrained clinical decision-making, given the limited data available per subject (4).

NMMs address part of this challenge by coarse-graining (e.g., mean-field approximation) the activity of neuronal subpopulations (e.g., pyramidal cells, excitatory/inhibitory interneurons of a cortical column) into a small number of state variables, typically interpreted as average membrane potentials or firing rates. Classical NMMs of coupled cortical columns and thalamo-cortical circuits have shown that such mesoscopic descriptions can reproduce a wide range of spontaneous and evoked electrophysiological phenomena, including rhythmic activity and visual evoked potentials (5, 6). In the context of epilepsy, NMMs form the basis of the Virtual Epileptic Patient framework, where they have demonstrated clinical value in localizing the epileptogenic zone (7, 8) and in comparing candidate surgical resection strategies *in silico* (9, 10). Model-derived estimates have been shown to correlate with post-surgical seizure outcomes (8, 11) and are currently undergoing prospective clinical evaluation (EPINOV, NCT03643016). Because they can be directly linked to macroscopic observables such as electro- and magnetoencephalography (EEG, MEG), NMMs offer an attractive compromise between biophysical interpretability and computational tractability. In principle, they provide a natural relationship between structural connectivity (e.g., from diffusion MRI), functional constraints (e.g., from EEG/MEG), and neuromodulation protocols. Potential applications include the optimization of brain stimulation approaches and parameters, as well as personalized treatment planning.

However, when NMMs are scaled to (surface-based) whole-brain networks with realistic connectivity, delays, and region-specific parameters, the computational demands again become substantial. Typical workflows include fitting model parameters to patient data for personalization, performing sensitivity analyses, and evaluating alternative stimulation strategies, and optimizing stimulation configurations. These tasks involve large numbers of simulations, for example for gradient-based or sampling-based parameter inference or optimization. Many existing NMM libraries were not designed with such use cases in mind: they often rely on CPU-based numerical integration, limited vectorization, and bespoke implementations that are difficult to differentiate through or to integrate into modern machine learning pipelines. As a consequence, the community lacks frameworks that are simultaneously biophysically grounded, scalable to large networks, and natively compatible with differentiable programming.

In parallel, the field of artificial intelligence (AI) has advanced rapidly through a tight coupling of model design, numerical libraries, and specialized hardware. The success of deep neural networks on large-scale perception tasks was enabled not only by architectural innovations but also by the availability of GPU-accelerated, tensor-centric frameworks that made it straightforward to express models and automatically compute gradients at scale (12, 13). Libraries such as JAX extend this paradigm by combining a NumPy-like interface with just-in-time compilation, automatic vectorization, and automatic differentiation, targeting CPUs, GPUs, and TPUs from a single high-level code base (14). Neural mass whole-brain models share key mathematical structures with recurrent and convolutional neural networks, including recurrence, convolution-like spatial coupling, time-series dynamics, delays, and nonlinear activation functions. Moreover, AI model training and NMM parameter inference share related optimization requirements, including the need for automatic differentiation, backpropagation, and efficient largescale parameter exploration. These parallels suggest that design principles and technologies developed for modern AI can be leveraged to build the next generation of large-scale brain modeling tools.

In this work, we introduce *BraiNN*, a NumPy-like, Python-based framework for neural mass modeling built on top of JAX (14). *BraiNN* is designed to express whole-brain NMMs in a concise, array-oriented syntax while transparently exploiting XLA-based compilation and accelerator backends, achieving runtimes competitive with or superior to existing frameworks across both region- and surface-level simulations, including configurations with biologically realistic conduction delays. BraiNN delivers two primary capabilities: first, it enables fast, GPU-accelerated simulations of large-scale brain networks through a two-scale architecture— combining a region-level NMM network (Jansen–Rit (5) in this study) with a subject-specific cortical surface mesh— and biophysically grounded EEG forward modeling via reciprocity-based, personalizable lead fields; second, it provides a fully differentiable modeling stack integrated with a hybrid personalization pipeline that combines Bayesian optimization (BO) for global parameter exploration with gradient-based refinement for local convergence.

We start by introducing the equations of the Jansen–Rit NMM and of region- and surface-based whole-brain models, before describing the implementation of BraiNN and the personalization pipeline. Subsequently, we report how verification (including reproducing and refining a prior study (15) on the canonical synchronization and bifurcation dynamics of Jansen–Rit networks) and benchmarking against reference and state-of-the-art libraries were performed. We quantify the achieved acceleration and demonstrate, using demanding application examples, that BraiNN enables successful model parameter inference and the exploration of large parameter spaces with clinically acceptable computational resources and wall-clock time. We discuss the use of gradient-based and Bayesian optimization for personalized modeling. The goal is to introduce a modern, accelerator-aware library for whole-brain dynamics and EEG signal modeling that is both efficient at scale and directly usable for subject-specific personalization, and is planned for open-source release in the near future.

## Methods

### Neural mass model

For the present study, each network node is described by a Jansen–Rit NMM, which captures the mean-field dynamics of a cortical column comprising a population of pyramidal projection neurons receiving feedback from excitatory and inhibitory interneuron populations (5). The model combines a linear second-order synaptic operator that maps mean incoming firing rates to postsynaptic potentials (PSPs), and a static nonlinear sigmoid that maps membrane potentials back to firing rates.

For each node, the local state is described by a six-dimensional vector **y** = (*y*_0_, *y*_1_, *y*_2_, *y*_3_, *y*_4_, *y*_5_)^⊤^ ∈ ℝ^6^, where *y*_0_, *y*_1_, and *y*_2_ are the average membrane potentials of the pyramidal projection, excitatory interneuron, and inhibitory interneuron populations, respectively, and 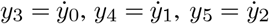 are their corresponding time derivatives. This augmented state formulation reduces the second-order synaptic dynamics to a first-order ODE system. Using the standard Jansen–Rit parameterization (5, 16), the local dynamics may be described as

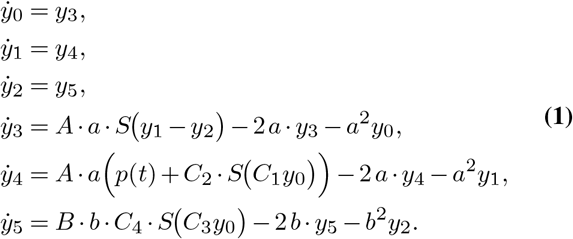

Here *A* and *B* are the average excitatory and inhibitory synaptic gains, *a* and *b* are inverse synaptic time constants, and *C*_1_, …, *C*_4_ are intra-column connectivity constants. The term *p*(*t*) represents the total excitatory input to the pyramidal population (external drive plus long-range coupling; see below). The firing-rate nonlinearity is given by

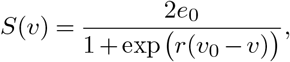

with *e*_0_ the maximum firing rate, *v*_0_ the firing threshold, and *r* the sigmoid slope. ‘Table 1 lists the default values and ranges. For the Jansen–Rit variant with stochastic noise, see Eq. (6).

**Table 1.** Parameter set for the Jansen–Rit neural mass model as originally proposed in (5) and widely adopted in subsequent studies (e.g., (16, 20)). The *Range* column consolidates bounds used in parameter-estimation studies (17) with broader simulator/model domains as implemented in The Virtual Brain (18, 19).

| Parameter | Default value | Range (17–19) |
| --- | --- | --- |
| $A$ | 3.25 mV | 2.6 mV to 9.75 mV |
| $B$ | 22 mV | 12 mV to 110 mV |
| $a$ | 100 s<sup>-1</sup> | 16.7 s<sup>-1</sup> to 500 s<sup>-1</sup> |
| $b$ | 50 s<sup>-1</sup> | 16.7 s<sup>-1</sup> to 500 s<sup>-1</sup> |
| $C_1$ | 135 | 32.5 to 2025 |
| $C_2$ | 108 | 26 to 1620 |
| $C_3$ | 33.75 | 8.125 to 506.25 |
| $C_4$ | 33.75 | 8.125 to 506.25 |
| $e_0$ | 2.5 s<sup>-1</sup> | 1.25 s<sup>-1</sup> to 3.75 s<sup>-1</sup> |
| $v_0$ | 5.56 mV | 3.12 mV to 6 mV |
| $r$ | 0.56 mV<sup>-1</sup> | 0.28 mV<sup>-1</sup> to 0.84 mV<sup>-1</sup> |

In the present study, the simulated local output voltage of a node is taken as the pyramidal PSP difference

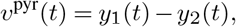

which is known to exhibit alpha-like oscillations and realistic evoked-response behavior for appropriate choices of *p*(*t*) and coupling parameters (5, 21).

### State representation and vectorization

Rather than implementing each node as an independent object, BraiNN aggregates all node states into joint tensors. For a network with *N* nodes we store the full state as a tuple of six arrays of size *N*. Furthermore, the right-hand side of Eq. (1) and all derived quantities are implemented as pure functions acting on these arrays. This design allows extensive use of JAX transformations such as `vmap` and `jit`, and enables compilation of the full network integration loop into a single accelerator kernel.

### Whole-brain region-level networks from structural connectivity

Whole-brain networks are constructed on a region-of-interest (ROI) parcellation. For each individual, we derive a structural connectivity matrix **W**_region_ ∈ ℝ^*N×N*^ and corresponding delay matrix **D** ∈ ℝ^*N×N*^ from diffusion-weighted MRI and T1-weighted MRI, following the image-based, personalized pipeline described by Karimi et al. (22). Briefly, a convolutional neural network-based segmentation algorithm jointly reconstructs MR images and segments up to 40 tissue classes from *k*-space data (23). The segmentation, together with diffusion tractography, is imported into Sim4Life (24) to generate subject-specific head models and white-matter connectivity matrices suitable for both electromagnetic and network simulations.

At the network level, each ROI is a Jansen–Rit node. Long-range coupling enters the pyramidal input term *p*_*i*_(*t*) of node *i* as

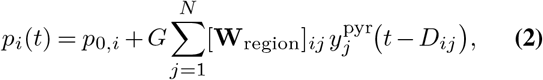

where *p*_*0,i*_ is the mean external drive, *G* is a scalar g coupling strength, and *D*_*ij*_ is the conduction delay between regions *j* and *i*. The delays are defined as the ratio of the inter-regional tract length–derived from diffusion tensor imaging–to a user-definable conduction speed, set to 3.9 mmms^−1^ in the present work. For nonzero *σ*_*ξ*_, this corresponds to a system of stochastic delay differential equations with distributed delays arising from the connectome (22, 25).

### Cortical surface representation and multi-scale mapping

BraiNN supports two simulation modes. In *region-level* simulations, each ROI corresponds to a single Jansen–Rit node coupled through **W**_region_ as described above. In *surface-level* simulations, the subject-specific cortical surface mesh (22–24) is used to assign a full Jansen–Rit node to each of its *M* vertices, providing within-region spatial detail that region-level simulations do not capture.

Each surface node *m* ∈ {1, …, *M*} is assigned to exactly one ROI *π*(*m*) ∈ {1, …, *N*} via nearest-neighbor mapping based on geodesic distance on the cortical surface. We denote by *S*_*i*_ = {*m* ∈ {1, …, *M*} | *π*(*m*) = *i*} the set of surface nodes belonging to ROI *i*.

In surface-level mode, each vertex *m* is governed by the full Jansen–Rit model Eq. (1), but its pyramidal input term receives contributions from two spatial scales. *Short-range* interactions between nearby surface nodes are mediated by a sparse geometric connectivity matrix **W**_surf_ ∈ ℝ^*M×M*^, which captures local lateral coupling on the cortical surface. *Long-range* coupling is inherited from the ROI-level structural connectivity: the outgoing signal of ROI *i* is defined as the average pyramidal PSP over its constituent surface nodes,

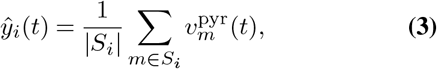

and all surface nodes within a receiving ROI share the same incoming long-range input. The full pyramidal input at vertex *m*, with *i* = *π*(*m*), is then

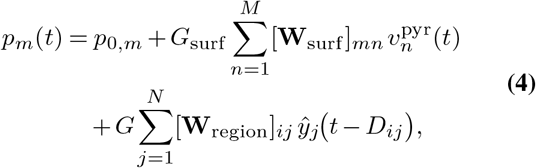

which parallels the region-level coupling (Eq. (2)) but adds the local surface term and replaces the region-level output 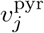 with the ROI-averaged surface signal *ŷ*_*j*_. The global coupling strength *G*_surf_ may differ from the region-level value *G*, reflecting the change in effective source density.

This two-scale architecture allows both classical region-level simulations (still widely used because of computational constraints), and surface-level simulations, which provide the spatial resolution needed for biophysically grounded EEG forward modeling via the lead-field projection described below.

Spiegler and Jirsa (26) showed that, for parcellations whose inter-node spacing falls within the spatial resolution of scalp EEG, a network of neural masses retains the spatial frequency content relevant to EEG-band signals with low attenuation. Surface-level simulations in BraiNN operate in this regime, offering a computationally efficient approximation that is appropriate when microscopic within-column detail is not required.

### EEG forward modeling via lead fields

To map simulated cortical activity to scalp EEG, we precompute a lead-field matrix 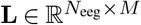 that linearly projects *M* cortical source strengths to *N*_eeg_ electrode voltages. Each cortical source is modelled as an equivalent current dipole oriented normal to the local cortical surface, so **L** is a scalar-per-vertex matrix obtained by projecting the full vector lead field onto the vertex normals. The lead field is computed using a reciprocity-based procedure (27–29): for each EEG electrode, a unit current is applied between that electrode and a common reference electrode and simulated using the electro-quasistatic finite-element solver (ohmic-current-dominated variant) with an image-based anatomical head model in Sim4Life. According to the Helmholtz reciprocity theorem, the normal components of the resulting electric field at the cortical vertices give the weights with which dipolar sources **d** located at these vertices and oriented normal to the surface contribute to the potential difference between that electrode and the reference. By assembling these weights in a matrix **L** (with rows corresponding to electrodes and columns to vertices), the simulated EEG is obtained as (27):

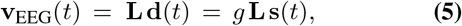

where **s**(*t*) is the vector of pyramidal-cell PSPs at each surface node (i.e., *v*_pyr,*m*_(*t*) in surface-level mode, or the ROI-broadcast values *v*_pyr,*π*(*m*)_(*t*) in region-level mode) and *g* is a factor that converts *v*_pyr_ (mV) to an equivalent current dipole moment (Am). This factor aggregates cortical column density, effective dipole length, and neuronal membrane surface area into a single scalar. Following standard practice in whole-brain network simulators (18), *g* is treated as a free parameter estimated during model personalization; when only relative EEG topography and spectral shape are of interest, *g* can be set to unity without loss of generality.

### Numerical implementation and acceleration

#### JAX/XLA-based implementation

BraiNN is implemented entirely in JAX, a NumPy-like array programming library that combines automatic differentiation with just-in-time compilation via XLA (14). All core components — the vector field, used to update the states of NMMs, network coupling, stochastic forcing, monitors, and objective functions — are expressed as pure functions over JAX arrays. This allows users to construct models using familiar NumPy-like code while seamlessly targeting CPUs, GPUs, or TPUs, abstracting away the burden of developing custom GPU kernels. Data interchange with existing scientific Python ecosystems is straightforward: NumPy arrays can be converted to JAX arrays and back with negligible overhead for typical problem sizes.

Crucially, the full trajectory integration is written as a single JAX `lax.scan` loop over time, wrapped in `jit`, so that the entire simulation (including monitors) is compiled into a single accelerator kernel. This ensures that all operations are vectorized over nodes. Furthermore, `vmap` can be used to further vectorize computation of trajectories on the same data, so that, for example, simulating multiple parameter sets or noise realizations can be simulated with only modest marginal cost.

#### Deterministic ODE solvers

For deterministic dynamics (noise switched off), BraiNN provides several explicit, nonstiff ODE solvers: forward Euler, Heun’s method (explicit trapezoidal rule), classical fourth-order Runge–Kutta, and adaptive embedded Runge–Kutta schemes of orders 5(4) such as Tsitouras’ TSIT5 and Dormand–Prince (Dopri5) (30, 31). All solvers are implemented in a solver-agnostic fashion, so that the same NMM/network definition can be integrated with different schemes without code duplication.

The Jansen–Rit equations normally have synaptic time constants on the order of at least 10 ms to 20 ms and moderate feedback gains for the parameter ranges considered in this work, providing a natural testbed for multi-scale integration schemes.

#### Stochastic differential equation solvers

To model endogenous fluctuations and measurement noise, BraiNN supports additive stochastic forcing of the form

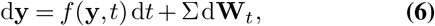

where *f* denotes the deterministic NMM/network right-hand side, Σ is a constant diffusion matrix, and **W**_*t*_ is a vector of independent Wiener processes. Since the noise is additive (i.e., independent of the state), the Itô and Stratonovich interpretations are equivalent (32). Therefore, we implement solvers for the additive-noise case, avoiding the additional complications of multiplicative or state-dependent noise.

Available SDE solvers include Euler–Maruyama and stochastic Heun schemes, as well as higher-order splitting Runge– Kutta methods for additive noise (32–34). These methods are again implemented in a fully vectorized and JIT-compiled fashion, allowing us to efficiently propagate large ensembles of stochastic trajectories for parameter exploration or uncertainty quantification.

#### Monitors and memory-efficient recording

BraiNN decouples the integration time step Δ*t* from the sampling interval Δ*t*_mon_ via a flexible monitoring interface. A monitor is a user-defined function that maps the full network state (and optionally derived quantities) at selected time points to a reduced set of observables (e.g., pyramidal PSPs, regional firing rates, surface-source activity, or EEG sensor voltages). Internally, monitors are evaluated within the `lax.scan` loop only at multiples of Δ*t*_mon_*/*Δ*t* (Δ*t*_mon_ is enforced to be a multiple of Δ*t*) and the resulting values are stored in preallocated arrays.

This design provides two benefits. First, it drastically reduces memory usage compared to storing the full state trajectory, which is particularly important for long simulations, large networks, or batched optimization runs. Second, it allows different observables to be recorded at problem-specific sampling rates without altering the underlying integration scheme. EEG time series, for example, can be sampled at 250 Hz to 1000 Hz, while internal fast variables are integrated at a much smaller Δ*t*.

#### Precision and parallelism

By default, simulations are executed in single-precision (32-bit) floating point to maximize throughput on modern GPUs. For verification and sensitivity analyses, BraiNN can be switched to 64-bit precision using JAX’s `jax_enable_x64` flag. All core components are compatible with `vmap` and `jit`, enabling parallel execution over batches of parameter sets, initial conditions, or noise realizations.

#### Implementation of delayed coupling

Conduction delays are implemented directly inside the compiled integration loop by storing the coupling variables in a fixed-size circular history buffer. At initialization, tract lengths are converted to integer delay indices according to

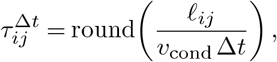

where *ℓ*_*ij*_ is the tract length from source region *j* to target region *i, v*_cond_ is the conduction speed, and Δ*t* is the integration step size. The buffer length is set by the largest resulting delay index, so memory scales with the maximum physiologically relevant delay rather than with the total simulation duration. During simulation, the current coupling output of each node, e.g., the pyramidal PSP, is written to the current buffer position and the write index is advanced modulo the buffer length. Whenever long-range input is evaluated, BraiNN gathers for all source–target pairs the entries at indices 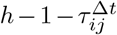, again modulo the buffer length, producing a tensor of delayed source activity that can be contracted with the structural connectivity matrix. This avoids explicit shifting of time-series arrays, keeps delayed coupling compatible with `jit`, `vmap`, and `lax.scan`, and allows delay-based network simulations to remain fully accelerator-executable.

### Personalization pipeline

#### EEG-based objective functions

To personalize model parameters for an individual subject, we define loss functions on EEG-derived features. Let *ϕ*(*·*) denote a feature-extraction operator that maps a multi-channel EEG time series to a summary feature vector, and let *ϕ*^⋆^ denote the corresponding target features computed from recorded data. Given simulated features *ϕ*(*θ*) from a model with parameters *θ*, we define the objective

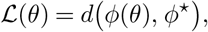

where *d* is an application-specific distance measure. In the present work, *ϕ* extracts the power spectral density (PSD) over 6 Hz to 30 Hz from each channel (35), and *d* is the Wasserstein distance between the target and simulated PSDs, providing a transport-based spectral mismatch metric that is robust to small frequency shifts between otherwise similar spectral profiles.

#### Gradient-based optimization in JAX/Optax

Because BraiNN is implemented in JAX, ℒ (*θ*) is, in principle, differentiable with respect to model parameters via automatic differentiation. We therefore expose the objective and its gradients to gradient-based optimizers from the Optax library — a JAX-native collection of gradient-processing and optimization routines (14, 36). In particular, we make use of the Adam optimizer (37) and variants thereof for fine-tuning parameters once a promising region of parameter space has been identified.

However, the combination of delayed recurrent coupling through the connectome and stochastic dynamics yields an effective recurrent neural network (RNN) with long temporal dependencies. It is well known that gradient descent in such systems is hampered by vanishing and exploding gradients and highly non-convex loss landscapes (38, 39). Moreover, the Jansen–Rit model exhibits rich bifurcation structure, including Hopf and saddle-node bifurcations, so that small parameter changes can induce transitions between qualitatively different dynamical regimes (16, 21). Together, these factors produce noisy, multi-modal optimization landscapes in which purely gradient-based methods readily become trapped in local minima or suffer from unstable gradient estimates, especially for long simulation windows.

#### Bayesian optimization with Gaussian process surrogates

To mitigate these issues and enable robust global exploration, BraiNN implements a complementary optimizer based on Bayesian optimization (BO) with Gaussian process (GP) surrogates (40, 41). In this setting, the unknown objective ℒ (*θ*) is modeled as a GP with mean function *µ*(*θ*) and covariance kernel *k*(*θ, θ*^*′*^). In particular, we use a Matérn-5*/*2 kernel together with a homoskedastic white-noise observation model to account for stochasticity in the simulation-based evaluations. Noisy evaluations of obtained from stochastic simulations update the GP posterior, which explicitly represents both the current best estimate and its associated uncertainty. In practice, this BO module is used for parameter spaces of moderate dimensionality, with *θ* typically having dimension ≤ 15. Initial kernel hyperparameters are inferred from variogram estimates and subsequently refined by maximizing the GP marginal likelihood using our own JAX implementation of the bounded L-BFGS-B optimizer.

We employ acquisition functions from a family of classical and numerically stabilized BO criteria, including expected improvement (EI) (42), probability of improvement (PI) (43), lower confidence bound (LCB), corresponding to the minimization form of GP-UCB (44), logarithmic expected improvement (LogEI) (45), corrected expected improvement (CEI) for noisy observations (46), and a logarithmic corrected expected improvement variant (CLogEI) combining the CEI correction with a LogEI-style reformulation for improved numerical stability (45, 46). In the EI case, new parameter candidates are selected by maximizing

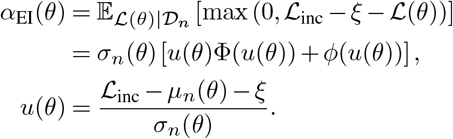

Here, *D*_*n*_ denotes the set of evaluations collected so far, and the expectation is taken with respect to the GP posterior predictive distribution of the latent objective,

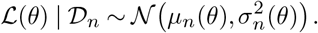

The functions Φ and *ϕ* denote the standard normal cumulative distribution function and probability density function, respectively. The parameter *ξ* ≥ 0 specifies the required improvement margin. In our implementation, the acquisition function returns the negative EI value because acquisition optimization is formulated as a minimization problem. In noisy settings, we take this incumbent to be the best current posterior estimate of the latent loss, rather than a single noisy simulation outcome. At each BO iteration, we first perform a global search over the parameter domain using Sobol sampling to identify promising candidate regions, and then refine the best candidate by locally optimizing the acquisition function with Adam. The selected parameter vector is subsequently evaluated by running a set of BraiNN simulations, often batched in parallel, and the resulting observations are used to update the GP posterior.

All GP computations, marginal-likelihood optimization, and acquisition optimization are implemented in JAX, allowing the entire BO loop — including forward simulations, feature extraction, surrogate updates, and hyperparameter fitting — to run on the GPU without having to return control to the host between iterations.

In practice, we use BO to perform global exploration and coarse localization of promising parameter regions, followed by local refinement using gradient-based optimizers from Optax. After BO identifies a promising basin, we refine the candidate by optimizing the GP posterior mean, and then, continue with global optimization of the original objective. This hybrid strategy leverages the strengths of both approaches: BO is robust to non-convexity and noisy objectives, while gradient-based methods efficiently exploit local curvature when gradients are reliable.

### Verification and benchmarking

#### Numerical verification and qualitative behavior

We verified the correctness of BraiNN’s neural mass and network implementation in several steps. First, we compared single-node and small-network trajectories against an independent CPU-based reference implementation of the Jansen–Rit model in TVB (18), using identical local parameters, inputs, and integration time steps. For each configuration, we compared simulated time series, power spectra, and basic summary statistics, including the mean and variance of the pyramidal-population output.

Second, we assessed whether BraiNN reproduced established qualitative network dynamics reported by Kazemi and Jamali (15). For this validation experiment, we replicated their parameter-sweep protocol for a network of stochastically driven Jansen–Rit nodes. We used the same nominal local parameters and coupling scheme as in the reference study and simulated a two-dimensional grid of configurations comprising a 256 *×* 256 ensemble, with 128 independent repetitions per grid point. This resulted in approximately 8.4 million simulations. The sweep covered coupling strengths from 0 to 19.5 and stochastic input levels from 0 Hz to 330 Hz. Each configuration was simulated for 40 s of equilibration followed by 40 s of analysis time, using a fixed integration step size of 0.5 ms and a monitoring step size of 10 ms, following the protocol reported by Kazemi and Jamali (15).

For each parameter combination, we computed the long-time average network synchronization and the dominant peak frequency of the node-wise average pyramidal-population power spectral density (PSD). To assess sensitivity to stochastic fluctuations, we additionally computed the across-realization standard deviation of the synchronization order parameter at each point in parameter space. These quantities were then compared qualitatively and quantitatively with the synchronization and peak-frequency profiles reported in the reference study.

#### Performance benchmarking

We benchmarked BraiNN with respect to runtime and scalability as a function of (i) the number of regions of interest (ROIs), *N*, (ii) the number of surface nodes assigned to each ROI, (iii) the conduction speed, which directly modulates propagation delays, and (iv) the number of trajectories evaluated in parallel. The structural connectome, delay structure, and the region- and surface level coupling matrices were taken from (22), while the corresponding benchmark variables were modified according to the respective experiment. All benchmarks were performed with fixed numerical integration settings, using a simulation time step of Δ*t* = 0.5 ms and a monitoring interval of Δ*t*_monitor_ = 10 ms. Each simulation comprised an equilibration phase of 2 min followed by a recorded simulation phase of 2 min.

Benchmarks were executed on both multi-core CPUs and GPU. The test system consisted of an AMD Ryzen Threadripper 3970X 32-core processor with 256 GB DDR4 memory and an NVIDIA GeForce RTX 3090 GPU. For each benchmark condition, we generated 15 random problem instances and repeated each experiment 5 times in order to obtain more robust runtime statistics. In the surface-resolution benchmark, the maximum number of surface nodes was set to ~ 19,000 for GPU experiments, whereas CPU experiments were restricted to at most 4,500 surface nodes due to performance limitations (Fig. 5).

Runtime measurements include the full end-to-end cost of simulation execution, including XLA compilation of the simulation kernel for the respective model configuration and device backend. Thus, the reported timings reflect the actual wall-clock cost encountered in practice rather than separating one-time compilation overhead from execution of the compiled kernel. This is a conservative accounting, since compilation is a one-time cost that is amortized when many simulations are run, *e*.*g*., during optimization. Finally, we profiled the computational cost of personalization by measuring the runtime per optimization iteration.

Note on nomenclature. Throughout this manuscript, references to TVB denote the official open-source simulator The Virtual Brain (TVB) framework (18). Performance benchmarks were performed using the current reference implementation available through the TVB software repository (commonly referred to as tvb-root). When possible, we compared BraiNN’s performance with four other libraries which perform forward simulations at the whole-brain level. A comparison of their supported features can be found in table 2, where:

**Table 2.** Feature summary of whole-brain simulation libraries considered in the benchmarking comparison. *Surface* refers to cortical-mesh simulations with within-region spatial detail. *EEG-LF* denotes built-in lead-field-based EEG prediction, *Optim*. denotes support for parameter fitting or optimization, and *Sparse mon*. denotes support for spatially and/or temporally sparse monitoring. *Notes*. ∷ indicates partial support, typically through custom code, external workflows,or framework-level functionality rather than a dedicated built-in whole-brain feature. α TVB itself is primarily CPU-based; GPU acceleration has been explored in separate TVB-HPC extensions rather than being part of the standard TVB simulator. Although TVB-inspired alternatives with hardware acceleration exist, including vbjax, VBI (50), and Fast_TVB (51), these tools do not support surface-based simulations. We therefore selected vbjax as a representative example of this class of TVB alternatives.

| Library | Acceleration |  | Biological plausibility |  |  | Model fitting & observability |  |  |
| --- | --- | --- | --- | --- | --- | --- | --- | --- |
|  | GPU | TPU | Region | Delays | Surface | EEG-LF | Optim. | Sparse mon. |
| BraiNN | ✓ | ✓ | ✓ | ✓ | ✓ | ✓ | ✓ | ✓ |
| TVB (18) | ✗<sup>a</sup> | ✗ | ✓ | ✓ | ✓ | ✓ | ~ | ✓ |
| vbjax (47) | ✓ | ✓ | ✓ | ✓ | ✗ | ✗ | ✓ | ~ |
| cuBNM (48) | ✓ | ✗ | ✓ | ✓ | ✗ | ✗ | ✓ | ✗ |
| brainmass (49) | ✓ | ✓ | ✓ | ✗ | ✗ | ✗ | ✓ | ~ |

- *Acceleration* refers to what type of hardware acceleration the particular library supports;
- *Region* refers to simulations where only region-to-region (structural) connectivity is considered;
- *Delays* refers to simulations where inter-regional communication includes conduction delays (yielding biologically realistic propagation delays);
- *Surface* refers to simulations where a cortical mesh is used to provide finer-grained spatial detail of regional cortical topography. In particular, surface-based simulations are required for spatially detailed EEG/MEG topographies.

#### Model optimization and personalization

We evaluated BraiNN’s model personalization capabilities via parameter-recovery experiments on synthetic data. In each experiment, a ground-truth parameter set was constructed from the default model ranges by randomizing the fitted parameter vector

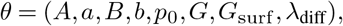

yielding an 8-dimensional optimization problem. The first four parameters correspond to Jansen–Rit model parameters listed in Table 1, while *p*_0_ denotes the mean external drive, *G* denotes the regional coupling scale, *G*_surf_ the scaling of surface coupling, and *λ*_diff_ the amplitude scaling of stochastic noise injected into the model. For each ground-truth parameter set, simulated EEG was generated from the corresponding network using a stochastic Heun simulator with an integration time step of 0.5 ms, a monitor sampling interval of 5 ms, and a 30 s recording window preceded by a 30 s equilibration period.

To recover the ground-truth parameters, we combined a PSD-based objective function, evaluated on 100 linearly spaced frequencies between 6 Hz to 30 Hz, with optimization of the parameter vector. For each ground-truth target, we generated 10 independent runs, each initialized with randomly sampled values for all eight fitted parameters and distinct random seeds for the injected stochastic noise. Both optimizers— global Bayesian optimization (BO) and standalone Adam optimization—were then applied to this same set of 10 runs under matched settings, enabling a controlled comparison between the two methods. We fixed the sequential optimization budget at 250 iterations for both fitting approaches to provide a fair, matched compute budget across methods while empirically ensuring convergence of the best-so-far loss. For each run, we quantified fit quality in terms of PSD similarity between simulated and target data (Wasserstein distance). Across the 10 runs, we further tracked the best-so-far loss over the course of optimization and computed its mean and standard deviation, characterizing both the typical fit quality and its variability across independent initializations. We additionally recorded the number of forward simulations and the wall-clock time for each optimization strategy.

## Results

### Numerical verification and qualitative behavior

BraiNN reproduced established Jansen–Rit dynamics at both the single-node and network levels. After validating single-node trajectories and small-network simulations against an independent reference implementation, we evaluated whether BraiNN could reproduce the qualitative synchronization behavior previously reported for stochastically driven Jansen–Rit networks by Kazemi and Jamali (15).

The synchronization maps obtained with `BraiNN` reproduce the qualitative structure reported by Kazemi and Jamali (15). In particular, the simulations show a clear coupling-dependent transition from weakly synchronized activity at low coupling strengths to strongly synchronized network dynamics at higher coupling strengths. This agreement supports the correctness of the implemented Jansen–Rit node dynamics and of the network coupling scheme.

The peak-frequency profiles also follow the expected qualitative behavior across the explored parameter space. Together with the synchronization maps, these results indicate that `BraiNN` captures the main dynamical regimes of stochastically driven Jansen–Rit networks. Small quantitative differences with respect to the reference results are consistent with the methodological limitation noted above, namely that the original study does not fully specify all stochastic-input implementation details. Importantly, these differences do not alter the main qualitative agreement between the reproduced and reference dynamical regimes.

In addition to the mean synchronization and peak-frequency maps, we quantified the across-realization variability of the synchronization order parameter. The resulting variability maps reveal a band of elevated fluctuations that are characteristic of the transition from weakly synchronized to highly synchronized dynamics. This pattern is consistent with increased sensitivity close to bifurcation-related regimes and with the complex dynamical behavior reported by Kazemi and Jamali (15). Overall, the replication demon-strates that `BraiNN` reproduces the expected qualitative network dynamics while additionally identifying the parameter regions in which the simulated dynamics are most sensitive to stochastic fluctuations.

#### Performance and scalability

We benchmarked BraiNN across region-level, surface-level, delay-coupled, and batched simulation configurations, comparing GPU-accelerated runs on an NVIDIA RTX 3090 to CPU baselines on a 32-core AMD Ryzen Threadripper 3970X.

For region-level simulations (Fig. 4), BraiNN on GPU achieved runtimes of approximately 5.5 s across the full range of *N* = 10 to *N* = 250 regions, representing a speedup of roughly 10.7 × over BraiNN on CPU and 215 × over TVB at *N* = 250. Notably, GPU runtimes for BraiNN remained approximately constant with increasing *N*, indicating that the network sizes considered here remain well within the parallelism budget of the GPU.

**Fig. 1.**
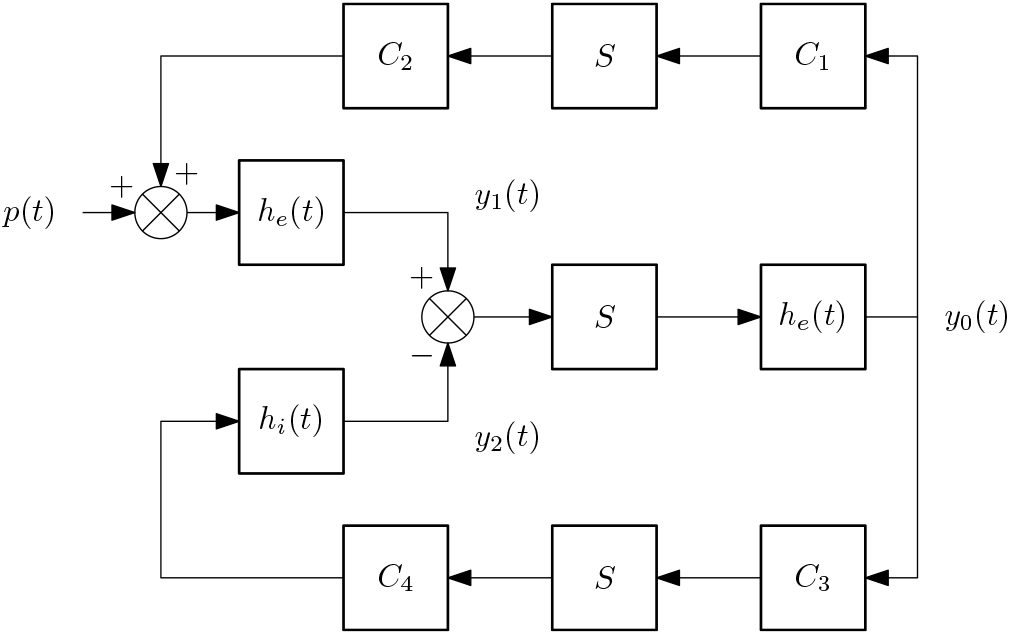
Schematic of voltage interactions in the Jansen–Rit model, showing how excitatory and inhibitory interneuron populations interact with the pyramidal cell population through feedback loops that shape the net membrane potential.

**Fig. 2.**
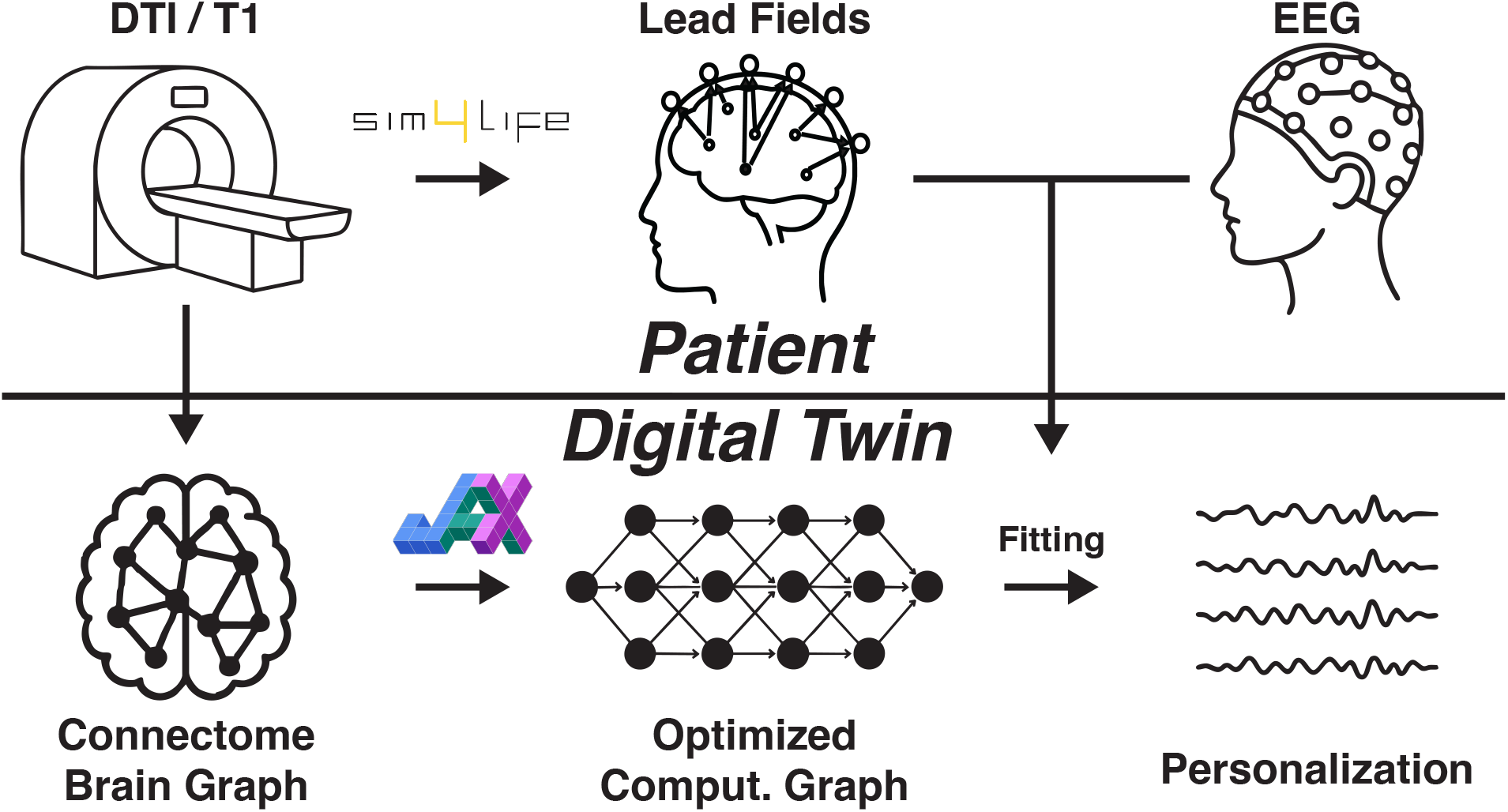
Conceptual overview of the patient-specific digital twin workflow implemented in `BraiNN`. Diffusion MRI/DTI data are used to construct a subject-specific structural brain graph based on its connectome, which is translated in `BraiNN` into an optimized, differentiable computational graph using JAX. Forward simulation of the resulting whole-brain model yields predicted EEG for the subject’s digital twin; model parameters are then fit by matching simulated EEG to the recorded EEG, enabling data-driven personalization of the digital twin.

**Fig. 3.**
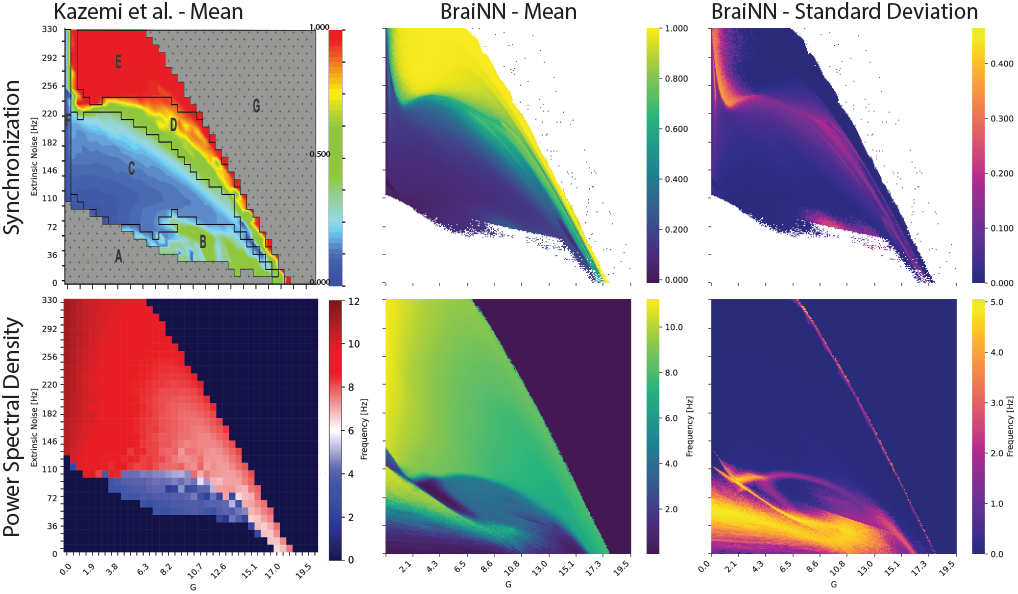
Coupling-dependent synchronization statistics for the parameter sweep used to reproduce the results of Kazemi and Jamali (15). From left to right, the columns show the reference synchronization values reported by Kazemi and Jamali, the mean phase synchronization obtained with `BraiNN`, and the corresponding standard deviation across realizations and conditions. Elevated synchronization variability highlights parameter regions with increased sensitivity and stronger bifurcation-related effects.

**Fig. 4.**
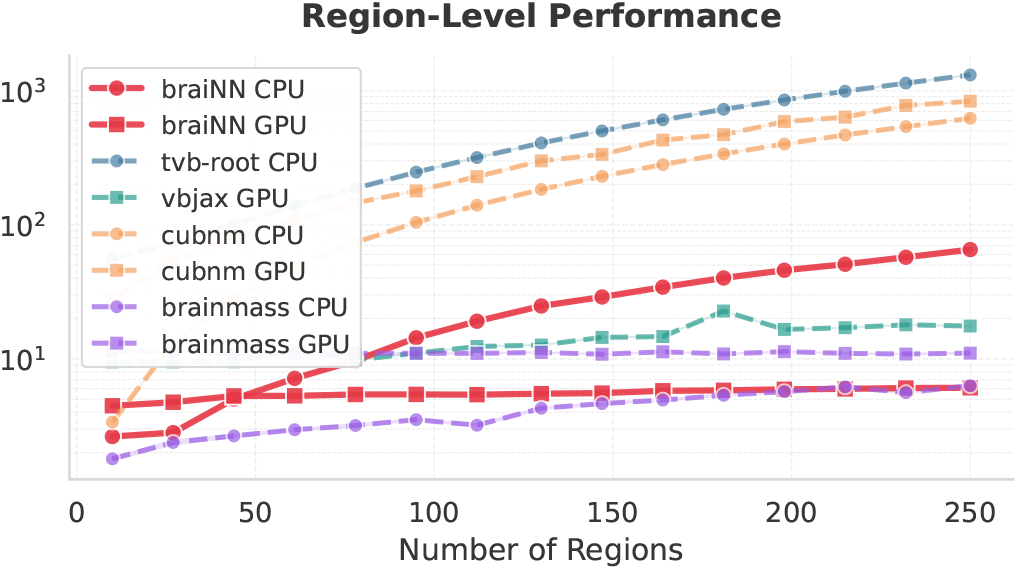
Runtime benchmarking for region-level simulations. Performance is reported as a function of the number of ROIs and compared across simulation libraries and execution backends as indicated in the legend.

For surface-level simulations (Fig. 5), BraiNN showed substantially better scaling than TVB. At the highest resolution benchmarked for both implementations, 4 392 surface vertices, TVB on CPU required approximately 1025.1 s, whereas BraiNN on GPU completed the simulation in approximately 12.9 s, yielding a speedup of approximately 79.5 *×*. BraiNN also supported substantially larger surface simulations, reaching 20 000 vertices in approximately 31.6 s. This corresponds to more than 4.5 *×* the largest available TVB surface benchmark while still running over 32 *×* faster than TVB at 4 392 vertices. Among the libraries evaluated, only BraiNN and TVB support surface-level simulations (Table 2).

**Fig. 5.**
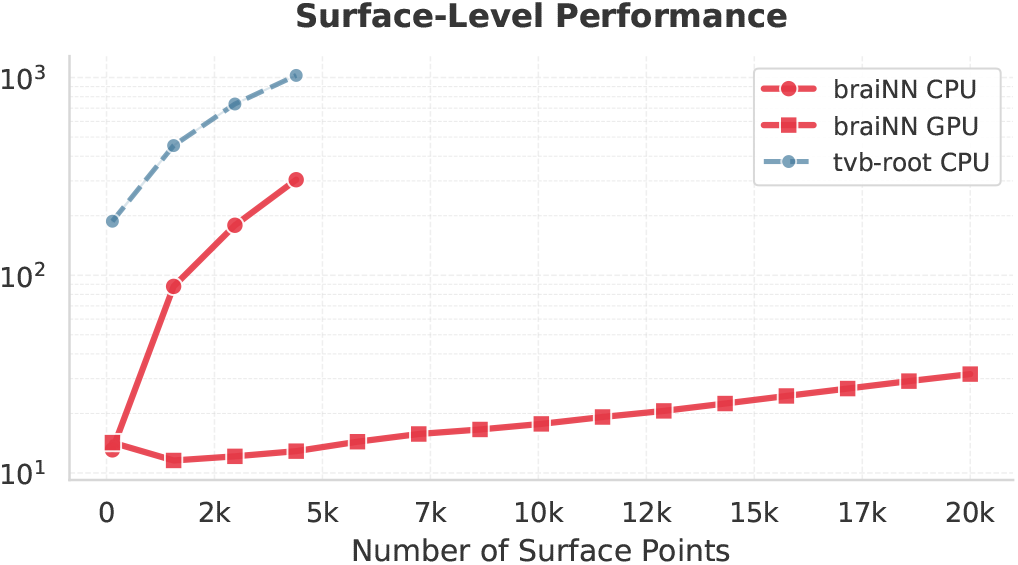
Runtime benchmarking for surface-level simulations. Performance is reported as a function of the number of surface points, and compared against TVB.

Increasing the maximum conduction delay (Fig. 6) increased runtimes for all libraries, as larger delays require longer history buffers. BraiNN on GPU showed only a small runtime increase, from approximately 5.0 s without delays to approximately 5.4 s at a maximum delay of 12.4 s, remaining competitive with or faster than the other GPU-accelerated implementations across the full delay range. BraiNN on CPU showed a similar delay-dependent trend and remained comparable to the CPU implementation of cuBNM across all tested maximum delays. This indicates that BraiNN’s delay handling introduces only modest overhead on both GPU and CPU, while preserving competitive performance relative to specialized GPU- and CPU-based simulators.

**Fig. 6.**
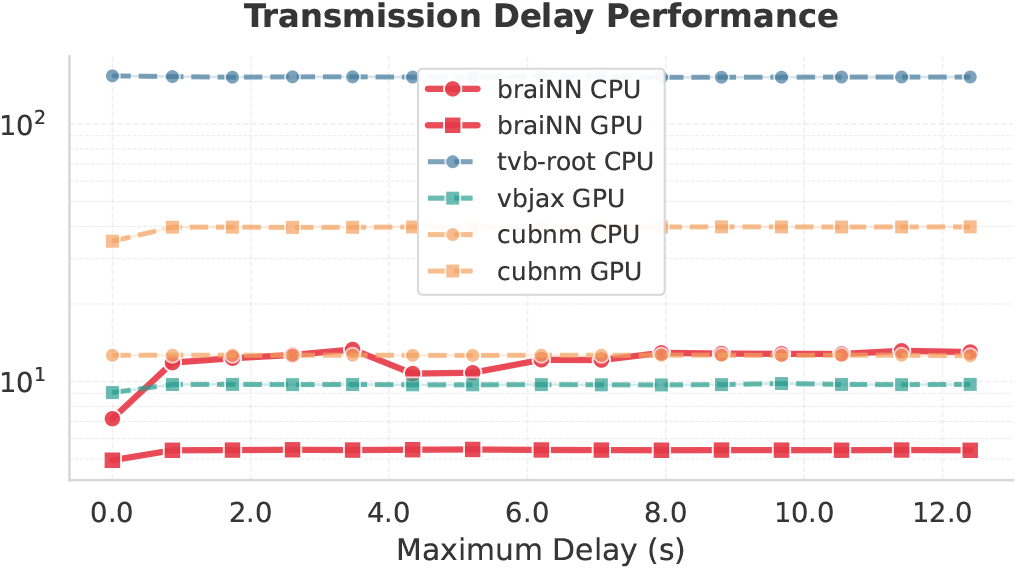
Runtime benchmarking as a function of the maximum conduction delay used in delayed coupling. Performance is compared across simulation libraries and execution backends as indicated in the legend.

Finally, batch evaluation (Fig. 7) demonstrated that GPU run-times for BraiNN scaled sublinearly with the number of parallel trajectories, reaching only 6.0 s for 32 simultaneous simulations compared to 4.6 s for a single run. This near-constant marginal cost per additional trajectory is a direct consequence of the `vmap`-based vectorization strategy described above, and is particularly relevant for the batched evaluations required during BO.

**Fig. 7.**
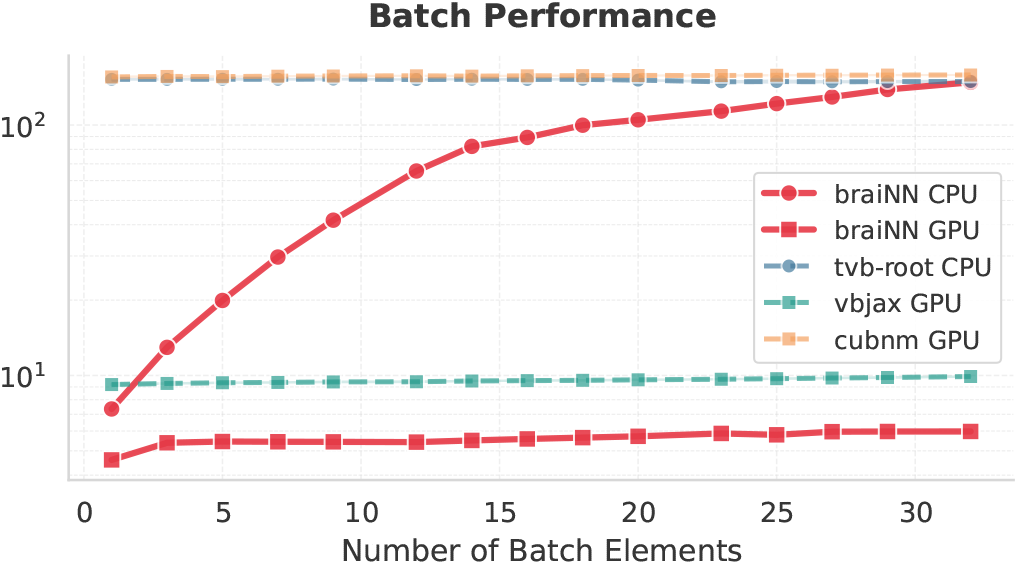
Total runtime benchmarking as a function of batch size (number of simulations evaluated in parallel). Performance is compared across simulation libraries and execution backends as indicated in the legend.

#### Parameter inference

We assessed BraiNN’s personalization performance by comparing the PSD of synthetic data from a known reference configuration (ground truth) to the mean best-fitting PSD obtained by optimizing the eight-dimensional parameter vector. For each target, both optimizers—global Bayesian optimization (BO) and Adam-based gradient optimization—were applied to the same set of 10 independent runs, each initialized with randomly sampled values for all eight fitted parameters and a distinct seed for the stochastic noise. Over the course of optimization, we tracked the best-so-far loss and computed its mean and standard deviation across the 10 runs, characterizing both fit quality and run-to-run variability.

#### PSD recovery

The optimization trajectories and the resulting spectral agreement are shown in Fig. 8. From an initial spectral mismatch metric of ~ 0.08, BO reached a best-so-far loss of 0.029 *±* 0.002, and its average fitted PSD closely tracks the ground truth across the 6 Hz to 30 Hz band with little run-to-run spread. Stand-alone Adam optimization stagnated at 0.055 *±* 0.027—roughly double the Wasserstein distance and more than an order of magnitude more variable— and its fitted PSD remained visibly mismatched to the target. This is consistent with the noisy, non-convex loss landscape and the vanishing and exploding gradients expected for long, delayed, stochastic simulations, thus confirming the rationale for using BO for global exploration rather than gradient descent alone.

**Fig. 8.**
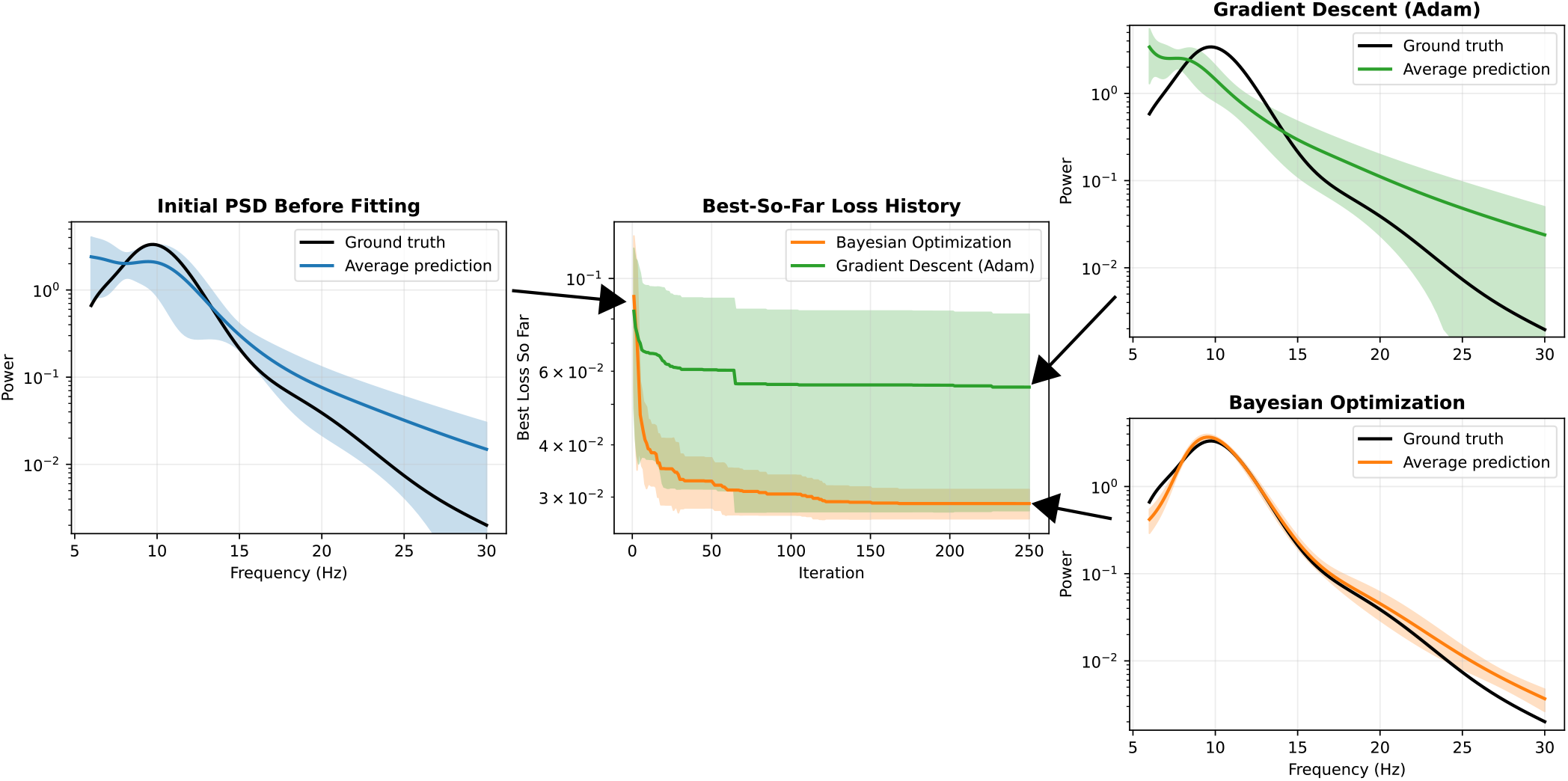
Parameter-recovery experiment comparing the two optimization methods in BraiNN’s personalization pipeline—global Bayesian optimization (BO) and stand-alone gradient descent (Adam)—on an eight-dimensional spectral-fitting problem by fitting the mean of 10 random initializations per method to a synthetic reference. *Left:* power spectral densities (PSDs) before fitting (reference and average over the initializations). *Middle:* best-so-far (running-minimum) spectral loss during fitting for both methods. *Right, top/bottom:* mean fitted PSD for Adam and for BO, overlaid on the reference. Shaded bands denote *±*1 standard deviation across the 10 fits; power uses a shared logarithmic scale. BO recovers the target spectrum closely and consistently (best-so-far Wasserstein distance of 0.029 *±* 0.002), whereas gradient descent alone stagnates at double that distance with ~10-fold higher variability (0.055 *±* 0.027) and requires 1.9*×* the wall-clock time.

#### Optimizer performance

On the RTX 3090, a complete BO fit required 140 min *±* 1.7 min versus 266 min *±* 45 min for gradient descent—1.9 *×* faster. Each BO evaluation cost 28.9 s—essentially one forward simulation (96 % of the 30.2 s a stand-alone run takes)—confirming that cost is dominated by the number of simulations rather than the BO machinery. Each gradient step cost 63.9 s—2.25 *×* a forward run and 2.2 *×* the BO per-evaluation cost—due to the automatic-differentiation forward pass through the long, delayed stochastic trajectory.

## Discussion

This work set out to develop a modern, accelerator-aware library for neural mass modeling that is both efficient at scale and directly usable for subject-specific personalization. BraiNN is implemented entirely in JAX and treats whole-brain simulations, feature extraction, and optimization as parts of a single, differentiable computational graph. The results show that this design yields a simulator that reproduces established Jansen–Rit dynamics, outperforms existing libraries in runtime, and offers a richer feature set (surface-level, delays, stochastic noise, EEG signal mapping with subject-specific lead fields, gradient and Bayesian optimization, spatio-temporally sparse monitoring among other features) for personalized whole-brain modeling.

On the modeling side, BraiNN follows the standard Jansen– Rit formulation, ensuring compatibility with prior work on mesoscopic cortical dynamics (5, 16, 20). By replicating the stochastic network experiments of Kazemi and Jamali (15), including the onset of synchronization and the elevated variability around critical coupling, we confirmed that the implementation captures the expected bifurcation structure in networks of Jansen–Rit oscillators (Fig. 3). Because the reference study did not fully specify the noise model, the minor discrepancies observed in low-coupling regimes were expected. This benchmark gives confidence that researchers can use BraiNN as a alternative for established NMM simulators while benefiting from substantially improved performance.

At the whole-brain level, BraiNN combines a region-level Jansen–Rit network with a subject-specific cortical surface and a reciprocity-based EEG forward model. This two-scale architecture is aligned with current trends toward individualized structural and functional mapping (1, 2, 22–24). Mapping ROI activity onto a cortical mesh and projecting it to EEG sensors via precomputed lead fields (24, 27, 28) enables simulations that are both computationally tractable and biophysically grounded at a level suitable for clinical workflows.

### Performance

Within the landscape of existing whole-brain libraries, BraiNN complements and extends tools such as TVB and its high-performance variants (18, 50–52) as well as newer GPU-focused frameworks like cuBNM, vbjax, and brainmass (47–49). Benchmarking results indicate that BraiNN achieves competitive or superior runtimes on CPUs and GPUs while supporting region-level, delay-based, and surface-based simulations in a single JAX/XLA stack (Table 2). Notably, none of the other GPU-accelerated brain simulators we evaluated currently support surface-based simulations. Recent TVB-derived efforts such as `vbjax` (47), Virtual Brain Inference (VBI) (50), and Fast TVB (51, 52) exploit GPUs or HPC resources for region-level models but do not incorporate cortical surface coupling and thus cannot natively produce spatially detailed EEG/MEG signals. This combination of performance and feature richness is central to enabling rapid experimentation and large parameter sweeps in both research and translational contexts.

The personalization experiments illustrate the benefits and limitations of exploiting automatic differentiation in large-scale brain models. Expressing EEG-based objectives in JAX allows the use of gradient-based optimizers from Optax (36, 37), but delayed recurrent coupling, stochastic dynamics, and long simulations lead to highly non-convex, noisy loss landscapes with vanishing and exploding gradients (16, 21, 38, 39). The hybrid strategy adopted here—global exploration with GP-based BO followed by local gradient-based refinement—yields accurate spectral recovery on synthetic data at reasonable computational cost (35, 40, 41). Thanks to accelerator-enabled batch evaluations, personalization required only a few hours for a whole-brain model of nearly 19,000 nodes, a speed that, to our knowledge, has not previously been reported for this scale of biophysically grounded modeling.

### Clinical acceptability

For personalized brain modeling to inform clinical decision-making, the computational cost of model fitting must be compatible with clinical workflow timescales. In practice, neuromodulation treatment planning unfolds over days to weeks: imaging acquisition, head-model construction, electrode planning or coil positioning, and multidisciplinary clinical review typically span multiple sessions before stimulation begins (22, 53). Within this timeline, the rate-limiting computational step has traditionally been model personalization. Early TVB-based fitting efforts reported approximately one hour of CPU time to fit a single 1 s EEG epoch at the region level using random Monte Carlo search (54); scaling such approaches to longer recording windows, surface-level models, or gradient-free global optimizers compounds the cost further. More recent simulation-based inference (SBI) approaches, such as VBI (50), reduce the repeated computational burden by training a neural density estimator up-front, after which per-subject inference requires only seconds. However, this strategy entails a substantial one-time training cost tied to a fixed model class and parameter prior, requires repetition, for example, when the connectome changes, and has so far been applied only to region-level models.

BraiNN’s hybrid BO/gradient pipeline occupies a different point in this trade-off space: it requires no offline pre-training, operates on surface-level models with ~ 19000 nodes and biophysically grounded EEG forward modeling, and completes a full spectral-target fit in approximately 2.3 hours on a single consumer GPU (NVIDIA RTX 3090). This runtime is well within the multi-day clinical planning window described above and, crucially, does not require high-performance computing infrastructure. In the context of a realistic clinical pipeline (22, 55)—where upstream steps such as image acquisition, segmentation, and connectome extraction themselves require on the order of hours—a model personalization step of this duration does not represent a bottleneck. We therefore argue that the combination of surface-level fidelity, consumer-grade hardware requirements, and fitting times on the order of a few hours places BraiNN within the practical envelope for integration into clinical neuromodulation workflows.

These results open a path toward clinical applications. For example, in neuromodulation therapy planning, BraiNN could be used to compare patient-specific predictions of network responses across different stimulation strategies to guide treatment decisions. However, a crucial next step is systematic validation on *real-world* EEG and multimodal datasets. All personalization results reported here are based on synthetic data generated by the same model class, and on PSD fitting alone. While this is appropriate for controlled identifiability studies, real data introduce additional noise, artifacts, and potential model mismatch. Evaluating BraiNN on empirical datasets, including patient cohorts, will be essential to validate robustness, refine modeling assumptions, and quantify performance relative to established workflows such as TVB- and VBI-based pipelines (18, 50).

### Limitations

Despite its benefits, BraiNN also has limitations that point toward important directions for future work. Currently, the library implements the Jansen–Rit cortical column model, though extension to other NMMs is trivial. Extending BraiNN to support additional neural mass models (e.g., thalamic relay populations, meso-circuit models of basal ganglia, or more detailed cortical field models (26)) will increase its versatility. The hybrid Bayesian optimization and gradient-based pipeline presented here provides an initial framework for parameter estimation; further development could incorporate more expressive surrogate models, amortized inference strategies, or tighter integration with proba-bilistic programming to improve scalability and uncertainty quantification.

On the feature-extraction side, the current objective relies on static power spectral densities (PSDs), but alternative feature choices may be more informative for predicting stimulation-induced changes in brain dynamics. Identifying objective functions better suited for fitting neural mass models remains a major area of ongoing research and offers substantial potential for future improvement. In particular, incorporating time-resolved or connectivity-based features such as dynamic functional connectivity (56) could improve sensitivity to transient network states and richer dynamical structure. Finally, while BraiNN’s current focus is on forward modeling and open-loop optimization, an exciting future direction will be closed-loop simulations for neuromodulation— embedding BraiNN into feedback control frameworks where stimulation parameters are adjusted online based on simulated neural responses.

## Conclusion

BraiNN provides a modern, JAX-based framework for neural mass modeling that bridges biophysically interpretable whole-brain models and the practical requirements of efficient simulation and personalization. By integrating subject-specific connectivity and head models, region- and surface-level dynamics, reciprocity-based EEG forward modeling, and hardware-accelerated integration in a single differentiable stack, it offers both high performance and a rich feature set. Numerical verification experiments confirm the correct reproduction of canonical Jansen–Rit dynamics, and bench-marking shows favorable scaling with network size, surface resolution, and delays. Across region-level, surface-level, and delay-coupled configurations, BraiNN achieves speedups of up to three orders of magnitude over established CPU-based frameworks such as TVB.

The tight integration with the JAX ecosystem enables flexible, differentiable EEG-based objectives and advanced optimization, including BO and gradient-based refinement. This combination supports efficient exploration of high-dimensional parameter spaces and opens a path toward individualized digital twins that can be tuned within clinically relevant time frames (22, 35, 40, 41). In synthetic parameter-recovery experiments, the full spectral-target personalization of an eight-dimensional parameter set for a surface-level model with approximately ~ 19,000 coupled nodes completes in a few hours on a single consumer GPU (NVIDIA RTX 3090), compared to multiple days with conventional neural mass modeling software. By drastically lowering the computational barrier, BraiNN enables more iterative and exploratory research on brain network dynamics—researchers can now perform expansive parameter sweeps or optimization studies that were previously prohibitive due to runtime constraints, placing whole-brain model personalization within the practical envelope of clinical neuromodulation workflows (22, 55).

Future work will focus on two main fronts: first, applying BraiNN to real-world clinical and research datasets to validate its robustness under realistic conditions and to assess its added value in practical personalization tasks; second, extending the framework to additional neural mass models, richer stochastic dynamics, more expressive EEG-derived features (including connectivity and dynamic functional connectivity (56)), and tighter integration with probabilistic inference and control toolkits (22, 50). As an open-source library, BraiNN invites the community to build upon it, extending its model library, improving its algorithms, and ultimately advancing the state-of-the-art in personalized whole-brain network modeling.

By unifying biophysically interpretable NMMs with modern accelerator-aware, differentiable computing, BraiNN will help to bridge the gap between detailed computational neuroscience models and the practical requirements of individualized neuromodulation.

## Acknowledgments

This research was funded in part by the Swiss National Science Foundation (SNSF) grant no. 10003470 (project TIME), and by Auden Techno Corp.

